# Circulating Exosomal PCNA: A Biomarker for Early and Pan-Cancer Detection

**DOI:** 10.1101/2025.02.06.636863

**Authors:** Weibing Shi, Xiaojie Sun, Haisheng Zhou, Hongting Da, Hong Li, Chunlin Cai

## Abstract

Detecting multiple cancers with a single, non-invasive liquid biopsy test remains a challenge, as no universally shared cancer biomarker with clinical significance has been identified in bodily fluids. In this study, we identified proliferating cell nuclear antigen (PCNA), a well-established pathological cancer biomarker in tissue biopsies, as being secreted into peripheral bodily fluids within exosomes. Our research demonstrated that cultured cancer cells continuously release large quantities of exosomal PCNA into their surrounding media. Furthermore, we confirmed its presence in human serum. Most importantly, cancer patients exhibited significantly higher levels of exosomal PCNA in their sera compared to healthy individuals.

To assess its diagnostic potential, we evaluated the sensitivity and specificity of exosomal PCNA in a cohort of cancer patients and healthy controls. Receiver operating characteristic (ROC) analysis revealed a sensitivity of 77.2% and a specificity of 94.4%. These findings establish exosomal PCNA in peripheral bodily fluids as a promising universal biomarker for the early detection, diagnosis, and monitoring of multiple cancers through liquid biopsy.

## Introduction

The global incidence of various cancers has risen significantly in recent years. Majority cancers are diagnosed at advanced stages, missing the optimal treatment window and leading to poor prognoses. Expanding early cancer detection and screening efforts is crucial for reducing cancer-related mortality and improving survival rates. Liquid biopsy offers a promising, non-invasive approach for large scale multi-cancer detection and screening, significantly enhancing treatment outcomes (1-2). Recently, development of single-test based liquid biopsy capable of detecting multiple cancers are of great interest. Some emerging technologies, such as circulating tumor cells (CTCs), circulating tumor DNA (ctDNA), and cell-free DNA (cfDNA), have shown potential in multiple cancer detection, but these technologies rely on complex sequencing and data analysis platforms, are limited by sensitivity and specificity, as well as the need for expensive platforms, restricting its affordability and accessibility.

Exosomes, a subset of extracellular vesicles (EVs), hold great promise as novel biomarkers for liquid biopsy applications (3-5). Compared to circulating tumor cells (CTCs) and cell-free DNA (cfDNA), these small lipid bilayer vesicles (30-150 nm in diameter) offer several advantages, including active secretion by living cells, high abundance, and stability in peripheral circulation, making them well-suited for clinical applications. Recent studies have demonstrated that exosomes carry various bioactive molecules, such as nucleic acids, lipids, and proteins, which are enriched in body fluids and play key roles in physiological processes, including tumorigenesis and cancer progression (6). The isolation and characterization of specific exosomal cargoes have already been applied in cancer diagnostics. For example, the ExoDx Prostate Cancer test (ExosomeDx) detects PCA3 and ERG biomarkers for prostate cancer (7, 8). Despite growing interest in exosome research and its potential in liquid biopsy, the development of a single test for detecting multiple cancers remains challenging due to the limited understanding of shared cancer biomarkers in exosomes from human bodily fluids.

In this study, we investigated and screened cancer cell proliferation factors using immunoassays to identify potential exosomal biomarkers shared across multiple cancer types. We identified proliferating cell nuclear antigen (PCNA) as a protein abundantly expressed in exosomes derived from all cancer types examined. PCNA plays a critical role in DNA replication and repair, serving as a molecular platform at the replication fork and functioning exclusively in proliferating cells (9-12). It has long been recognized as a cancer marker in tissue biopsy applications due to its high nuclear presence in proliferating cells. Although PCNA has traditionally been considered a nuclear protein, recent studies (13, 14) have reported its cytoplasmic localization, though its significance remains unclear.

In this study, we discovered secreted forms of PCNA—exosomal PCNA—present in human peripheral bodily fluids, including serum and urine. We developed unique and highly sensitive methods to detect exosomal PCNA in these fluids. Furthermore, we found that exosomal PCNA levels were significantly elevated in multiple cohorts of cancer patients compared to healthy controls.

Our findings pave the way for developing a single-test, multi-cancer liquid biopsy tool that is suitable for large-scale cancer screening, offering a more affordable and accessible diagnostic option.

## Materials and methods

### Cell culture

The cancerous cells used in this study were purchased from ATCC (Manassas, VA). Cells were all routinely cultured in the optimal media containing 5-10% fetal bovine serum (FBS), 2 mm l-glutamine, 100 U/ml penicillin-streptomycin, maintaining in the constant temperature and humidity sterile incubator with 5% CO2 at 37°C. The media were replaced regularly, followed by subculture when 85-90% cells were confluent.

### Exosome extraction

To extract the exosomes from serum samples, we followed a protocol from exosome extraction kit for plasma, ExoQuick (SBI, Cat #EQULTRA-20A-1). Briefly, 250 µL of serum was mixed with 67 µL of ExoQuick solution and incubated for 30 minutes at 4°C followed by centrifugation at 3000× g for 10 minutes. After centrifugation the extracellular vesicles appear as a beige or white pellet at the bottom of the tube. The pellet was resuspended and the exosomes were further isolated by a pre-packed purification column.

The exosomes in the cultured media were extracted through ultra-high-speed differential centrifugation. Before the extraction of exosomes, the cells were washed with phosphate-buffered saline (PBS) for three times, then, cultured in the media with 5% exosome-free FBS (System Biosciences) for 24hr. The cultured media were collected and centrifuged at 300 g for 10 min to remove cells and cell debris. After filtering through the 0.45 μm filter membrane, about 2ml of media was centrifuged at 120,000 g for 2 hr. The supernatant was discarded, and the pellet was re-suspended in 50ml exosome-free PBS, and centrifuged at 120,000 g for another 2 h to remove the residual proteins. The amount of the recovered proteins was measured using the BCA protein assay kit. Exosomes were used as fresh preparations for immunoblotting or were conserved at −80°C for later use.

### Western Immunoblotting

Immunoblotting was performed using standard protocols. Cells or prepared exosomes were lysed in 10 volumes of lysis buffer (50 mm Tris-HCl, pH 8.0, 0.5% Triton X-100, 150 mm NaCl, 20 mM NaF, 2 mM EGTA, 5mm EDTA, 1mm Na3VO4, 1 mm phenylmethylsulfonyl fluoride, 10 µg/ml Aprotinin/leupeptin/pepstatin), and protein concentrations were determined by BCA assay. Equal amounts of protein were separated by SDS-PAGE on 10–15% polyacrylamide gels and transferred to PVDF membrane (Millipore). The membranes were blocked by incubating with blocking buffer 1XTBST (50 mM Tris–HCl, pH 7.6, 150 mM NaCl, 0.1% Tween 20) with 5% skim milk for 1 h at room temperature. The blots were then incubated overnight with the primary antibodies in blocking buffer with gentle rocking at 4°C. The primary antibodies used are as follows: antibodies against PCNA (1:2000, Cell Signaling Technology/CST, #2586), MCM2 (1:2000, CST, #12079), E2F1 (1:1000, CST, #3742), CDK1 (1:2000, abcam, #ab131450), AURKA (1:2000, CST, #1475), PLK1 (1:1000, abcam, #17057), TROP2 (1:2000, CST, #194956). The blots were washed twice in blocking buffer at room temperature and incubated with horseradish peroxidase conjugated secondary antibody (1:5000 v/v) at room temperature for 1 h, then twice washing at room temperature. Immunoreactive bands were detected using the ECL Plus chemiluminescent detection system (T-4600, Tanon).

### Immunoprecipitation of exosomal PCNA

Immunoprecipitations of exosomal PCNA were performed as follows. Briefly, exosomes were lysed in TNE buffer (50 mM Tris-HCl pH 8.0 containing 150 mM NaCl, 5 mM EDTA, 1% Nonidet P-40, 5 mM NaF, 1 mM Na-pyrophosphate, 1 mM Na3VO4, 2 mM PMSF and 10 µg/ml aprotinin and leupeptin). Triton-X 100 was added to a final concentration of 1% and mixed for two hours at 4°C, followed by centrifugation at 20000 g for 20 min at 4°C. The supernatants were pre-cleared by incubation with GammaBind A/G-Sepharose (GE Healthcare), followed by overnight incubation with PCNA antibody (10 µg/ml extract) at 4°C. The immune complexes were harvested by incubation with GammaBind G-Sepharose (2 h, +4°C). After three 10 minute washes with TNE buffer and two five minute washes with PBS, the proteins were eluted in Laemmli’s sample buffer and resolved by SDS-PAGE.

### Enzyme-linked immunosorbent assay (ELISA) detection of serum and urinary PCNA

Serum PCNA was measured using ELISA kits obtained from Puer Biotechnology Co., Ltd. (Hefei, China) according to the manufacturer’s instructions. Briefly, after Leukocyte and Platelet-derived exosomes depleted using CD45 and CD61 antibody-coated magnetic beads, 50 μl serum samples were mixed with 50 μl 1XPBS containing 0.4% Tritron-X100 (to release PCNA from the exosome) for 30min at room temperature, then loaded onto ELISA plates precoated with monoclonal anti-PCNA and incubated at 37°C for 45 min. Subsequent to being washed four times with washing buffer, 50 μl/well anti-IgG conjugated with biotin was added and then the samples were incubated at 37°C for 30 min. The plates were washed again, streptavidin-HRP solution was added and the samples were incubated at 37°C for 30 min. Color developing agents A and B (50 μl/well) were added in sequence and incubated in the dark for 15 min. The reaction was terminated with stop buffer. Absorbance was measured at 450 nm on a microplate reader. The concentrations of PCNA were determined by interpolation from the standard curve.

### Patient samples

The clinical study was carried out according to the principles of the Declaration of Helsinki and was approved by the medical ethics committee of the First Affiliated Hospital of Anhui Chinese Traditional Medical University. Written informed consent was obtained from each subject. The plasma samples were collected from 161 healthy volunteers and 204 patients who had pathologically diagnosed with the following cancers including 26 with liver cancer, 20 with lung cancer, 26 with gastric cancer, 30 with colorectal cancer, 30 with breast cancer, 22 with ovarian cancer, 20 with pancreatic cancer, and 30 with prostate cancer. Of the cancer patients, there were 114 males (56%) with an average age of 65.08 ± 10.12 (range 35-88) years and 90 females (44%) with an average age of 63.27 ± 7.55 (range 46-82) years. Of the healthy volunteers, there were 85 males (53%) with an average age of 61.79 ± 11.52 (range 34-86) years and 76 females (47%) with an average age of 60.16 ± 6.09 (range 48-86) years.

### Statistical analysis

Statistics were performed using SPSS software, version 19.0 for windows (SPSS Inc, IL, USA). The biological and clinical characteristics of human subjects were shown as means ± SD (standard deviation). Two groups were compared with a Mann-Whitney test. The significance level was chosen to be P < 0.001.

## Results

### PCNA protein is present in the media of the cultured cancer cells

To explore the possibility that cell proliferation markers could be secreted and serve as molecular tools in cancer liquid biopsies, we used a dot scan immunoassay to analyze thirty-six cell proliferating markers in the cell media from three cultured cancer cells (HEK293, Hela, and COS7), and the presence of PCNA protein in the cultured media was identified. Next, PCNA along with six other commonly used proliferating markers (CDK1, AURKA, PLK1, MCM2, E2F1, TROP2) were selected for further testing. The presence of these proteins in the media of cultured HEK293, Hela, and COS7 cells was then examined by immunoblots with antibodies respectively. In these assays, cell lysates were included as the positive controls. PCNA protein was detected with a strong immunoactivity in both the cell lysates and the media of all three tested cells. In contrast, the other six proteins were only found in the cell lysates but absent in the media (**Figure 1**). To further confirm the presence of PCNA in the media of cultured cancer cells, two PCNA antibodies against the different epitopes were in-house produced. In SDS-PAGE analysis of the cell lysates and the cultured media, both of our PCNA antibodies confirmed the presence of PCNA in the lysate and medium of proliferating cells (HEK293, Hela, and COS7) in a direct immunoblotting or immunoprecipitation assay, but are absent in non-proliferating cells. These results suggested that PCNA protein is released extracellularly from the cancer cells.

**Figure 1.**
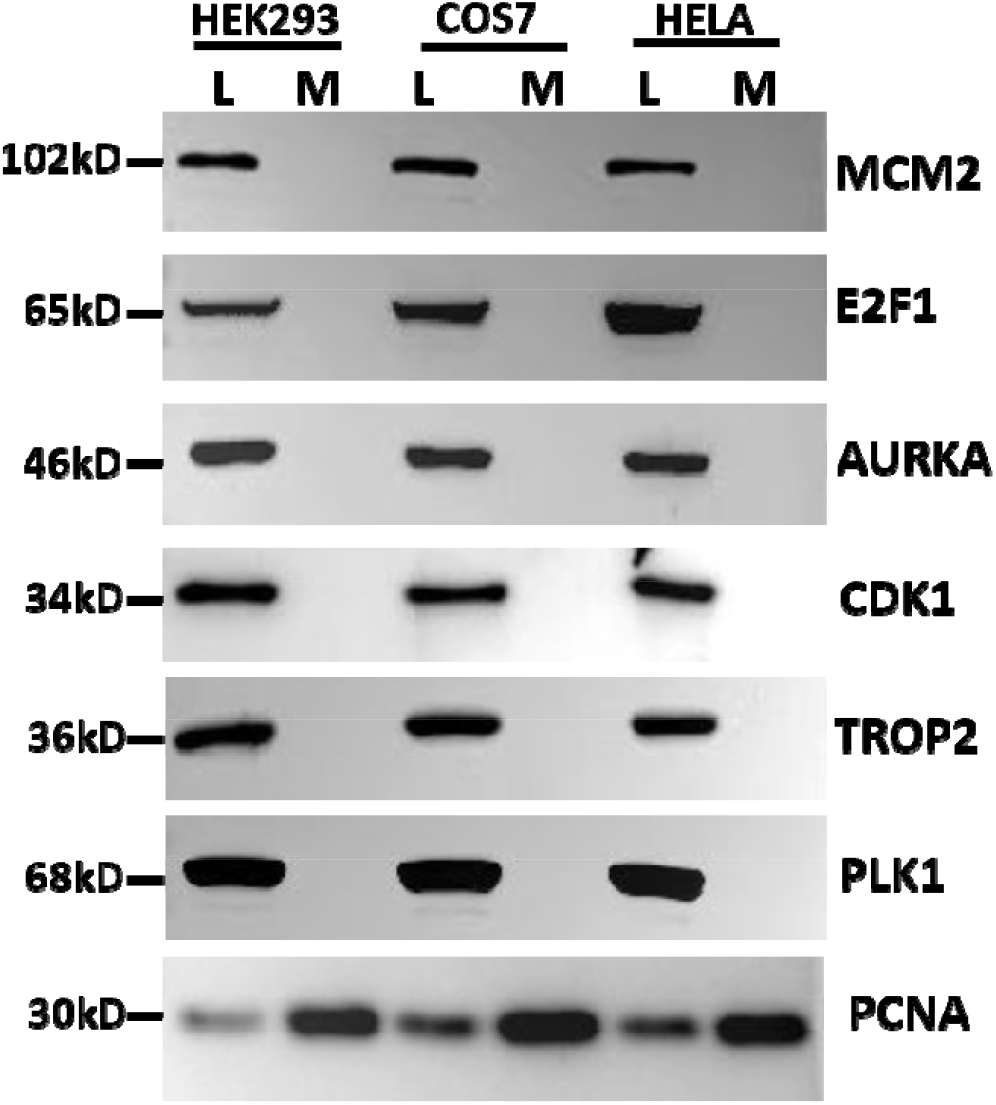
Immunoblot analysis of the presence of seven cell proliferation markers in three types of cultured cancer cells (HEK293, COS7, HeLa). L indicates the cell lysate, and M denotes the medium of the cultured cells.

### Cancer cells release PCNA protein via the exosomes

To investigate the presence of extracellular PCNA protein, we separated the substance in the culture media of the above cancer cells into 100kD-plus, 50-100kd, and 10-50kD fractions by PALL ultrafilters. PCNA protein is present in the fraction of 50-100kD, indicating it is not a free monomer in the cultured media. Based on the fraction range, we hypothesized that PCNA protein is present in the extracellular vesicles. We then purified exosomes from the cultured media of the cancer cells and found the presence of PCNA protein in the purified exosomes by immunoblots, no PCNA protein was detected in the media after the exosomes were depleted (**Figure 2**). Further, we studied tens of other different types of cancer cells and confirmed the presence of PCNA protein in extracellular exosomes in media of all cancer types examined (**Figure 2**). We also measured concentrations of the extracellular exosomal PCNA protein by ELISA and found it to have a 3.17%-5.24% of total exosomal proteins. Our findings demonstrated that PCNA protein is enriched in the exosomes secreted from cancer cells.

**Figure 2.**
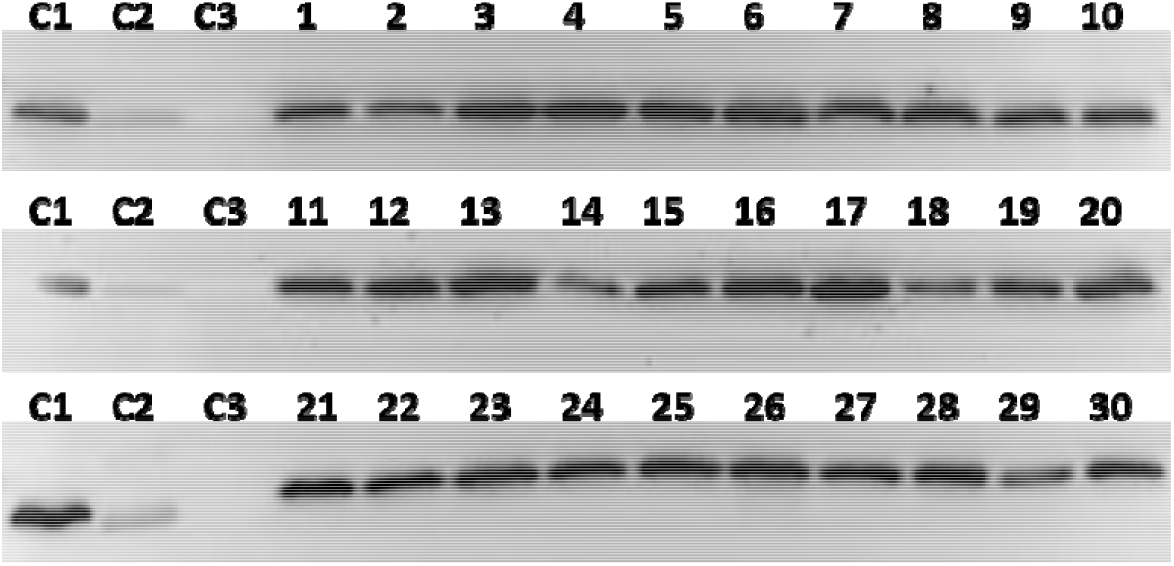
Immunoblot analysis of the exosomal PCNA levels in the media from 30 cultured cancer cells. C1-C3 indicate three controls for PCNA signal specificity comprising the lysate of HEK293 cells (C1), the lysate of embryonic mouse brain tissue (C2), and the lysate of mature mouse brain tissue (C3). Numbers 1-30 represent the following cancer cells: SMMC-7721(1), HepG2(2), Hep3B(3), Huh-7(4), MHCC97(5), A549(6), SKMES1(7), NCIH460(8), MGC803(9), SGC-7901(10), AGS(11), HSC-38(12), SGC-7801(13), HSC-66(14), MDA-MB-436(15), MDA-MB-435(16), BT474(17), MCF7(18), HCT116(19), SW480(20), HT29(21), Caco-2(22), SKOV3(23), A2780(24), ovcar3(25), PANC-1(26), CAPAN-1(27), PANC02(28), U87MG(29), SHSY5Y(30), respectively.

### PCNA protein is detected in serum of cancer patients

The discovery of PCNA protein in the media of cultured cancer cells was unexpected but reflects its potential clinical value in cancer liquid biopsies. Hence, we investigated whether exosomal PCNA is in the sera of cancer patients. We examined serum samples from a small pool of participants including 26 cancer patients and 26 normal controls. The exosomes from these subjects were purified, and immunoblotted with PCNA antibodies subsequently. Showed in **Figure 3**, PCNA protein was detected in all subjects. Markedly, cancer patients have a higher level of PCNA protein than healthy controls.

**Figure 3.**
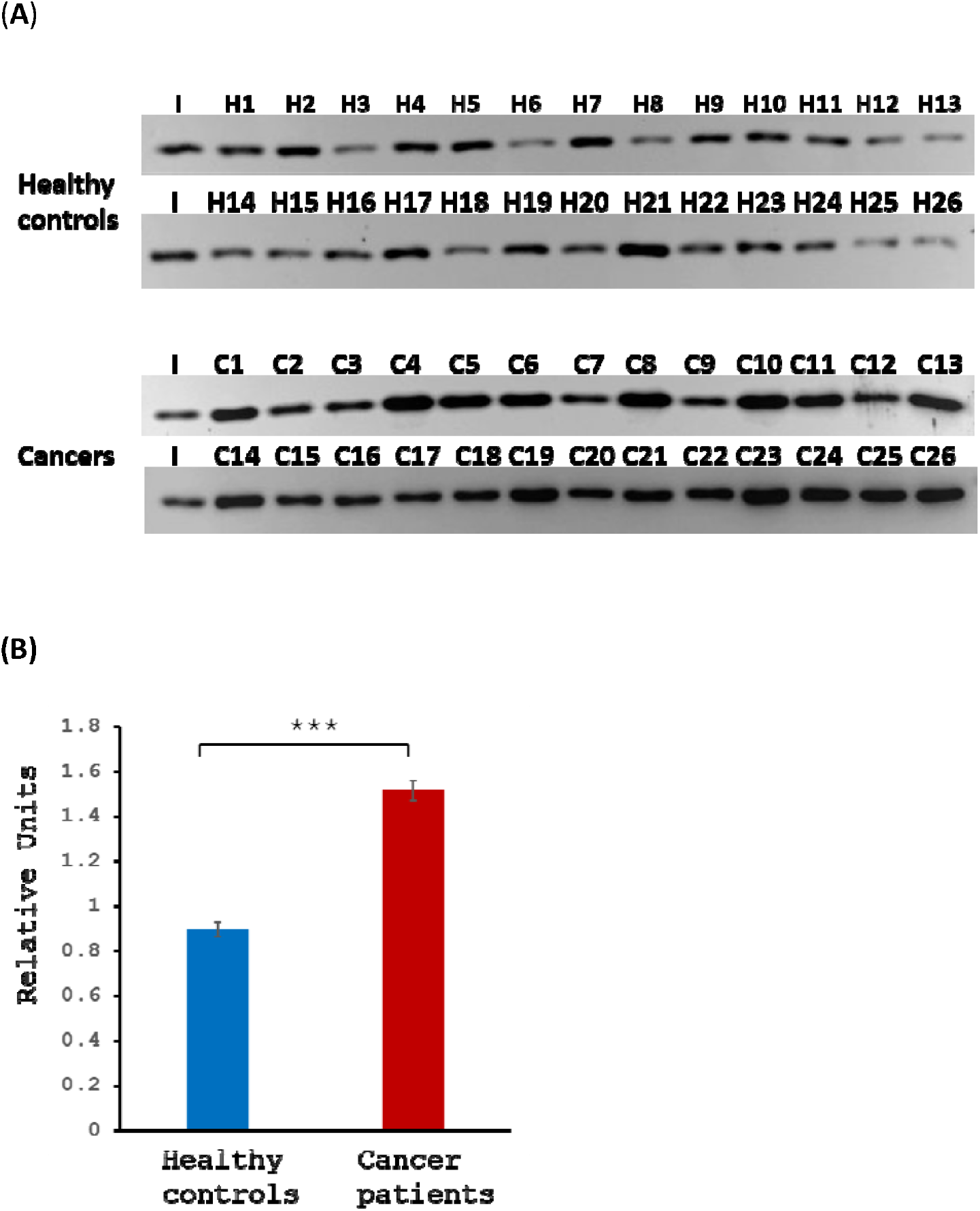
Immunoblot analysis of sera exosomal PCNA protein in healthy controls and cancer patients. (A) The two upper panels show the immunoblot analyses of sera from healthy controls (H1-H26) and the two lower panels from cancer patients (C1-C26). The sample I, indicated as an internal control, is a mixture of the sera from three healthy subjects. (B) Densitometric quantitation of the PCNA specific signal in (A).

### Exosomal PCNA protein is a potential serum biomarker for detecting and monitoring cancers

Our results show PCNA protein is present in human sera though it is a well-documented nucleus-located cell proliferating marker. We detected that exosomal PCNA protein is higher in the sera of cancer patients than in healthy controls. These results suggested the exosomal PCNA protein may serve as a molecular biomarker for cancer liquid biopsy for clinical applications, for example, early cancer detection before symptom appears, cancer diagnosing and progress monitoring, etc. We went on to further investigate the sensitivity and specificity of exosomal PCNA in detecting cancers by ELISA. A case-control study was also conducted. A total of 365 subjects were enrolled in this study, composed of 161 healthy controls and 204 cancer patients. ROC curves were plotted to define the optimal cutoff values of serum PCNA for cancer detection. Our study showed that with the optimal cutoff value of 97.81ng/ml in serum, PCNA had a sensitivity of 77.2%, specificity of 94.4%, and a positive predictive value of 45% for cancer diagnosis.

**Figure 4A.**
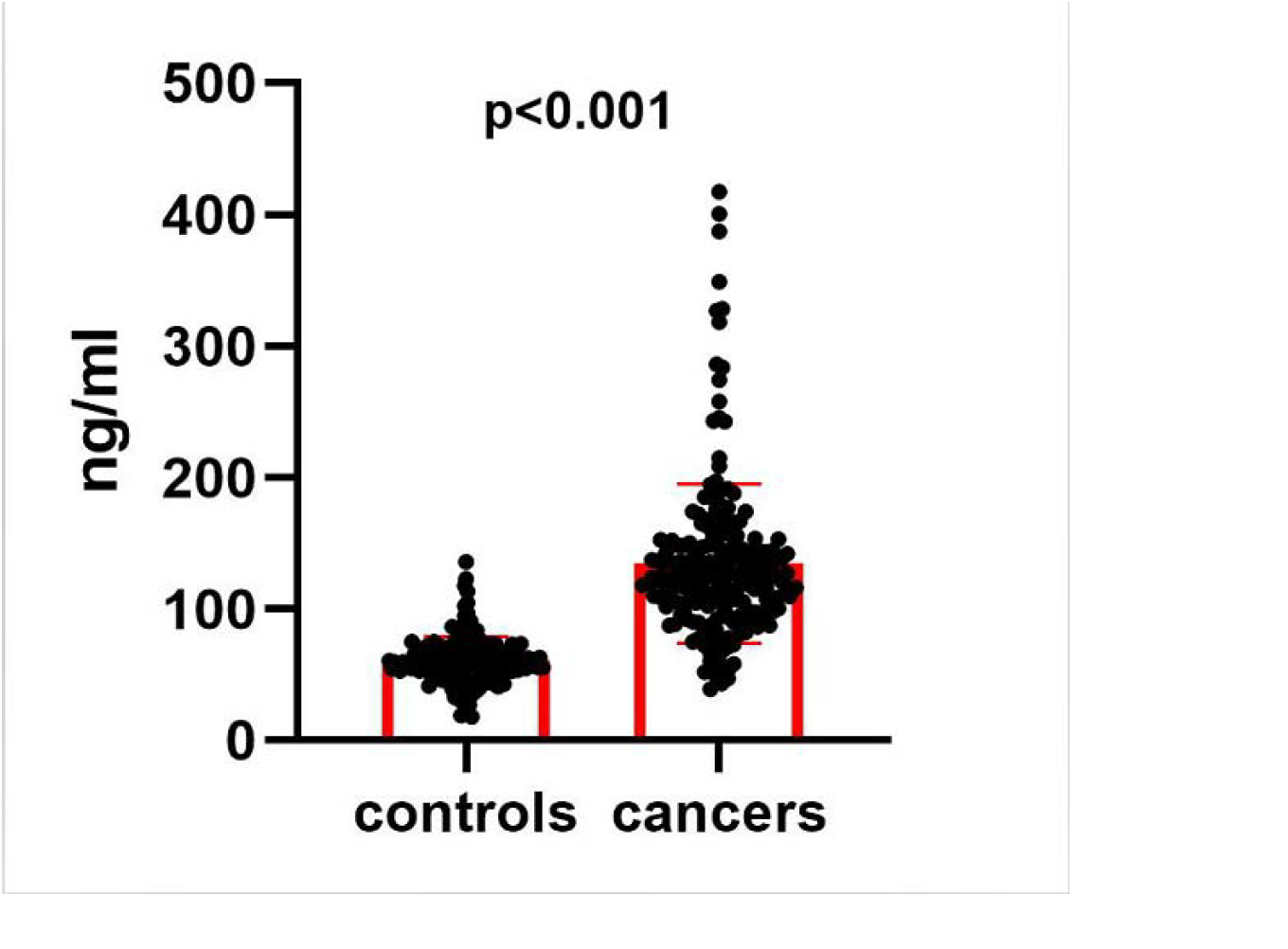
Scatterplot of serum PCNA levels in healthy controls and cancer patients detected by ELISA assays. The scatter dot represents a single value, and the horizontally thick line represents the mean. The horizontally light lines below and above the mean line represent respectively the mean minus and plus one standard deviation. Statistical significance (P<0.001) is indicated.

**Figure 4B.**
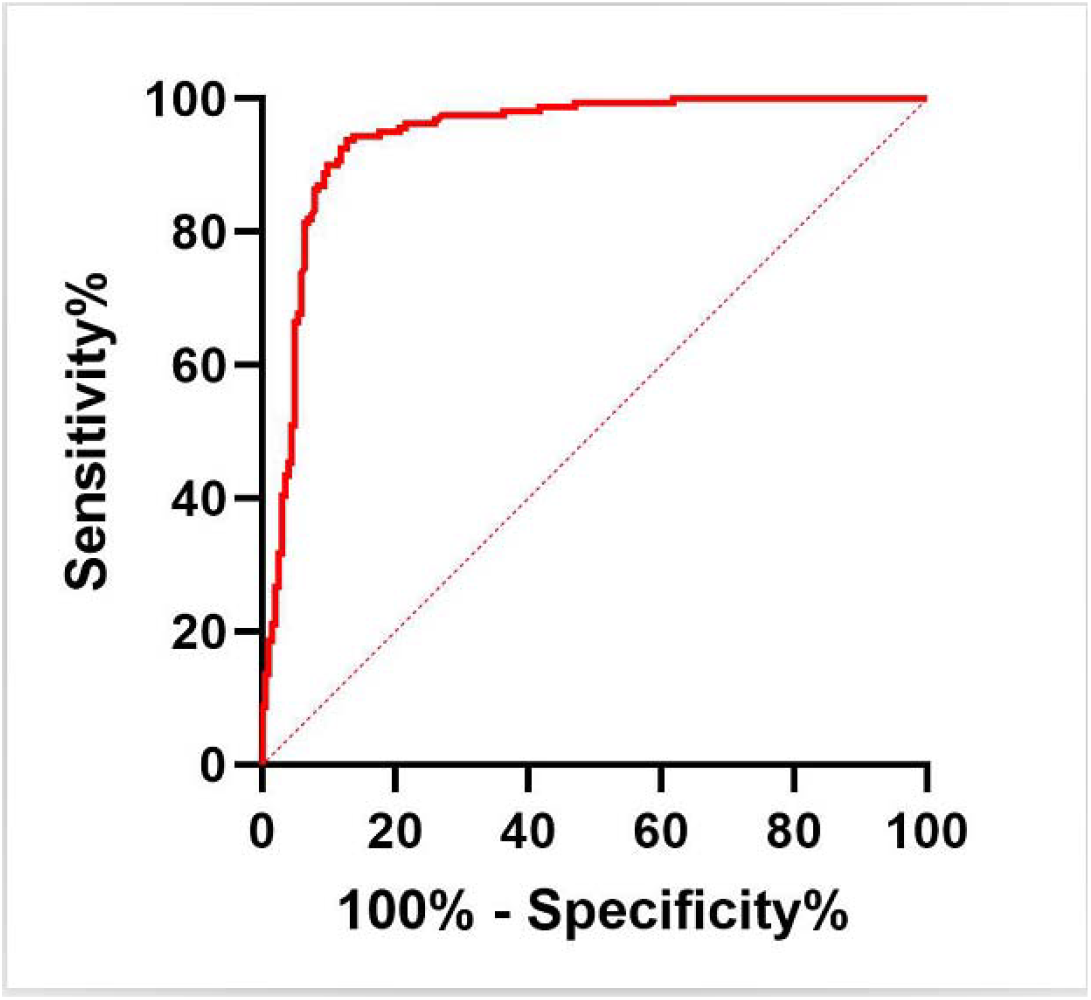
ROC curve of serum PCNA detected by ELISA assays for the diagnosis of cancers. The AUROC value for diagnosing cancers was 0.94 (95%CI: 0.92–0.97), with the sensitivity of 77.2% and the specificity of 94.4% under an optimal cut-off value of 97.2ng/ml.

## Discussion

PCNA protein has been generally recognized as a nuclear protein, and often as a histological biomarker of malignancies for multiple cancers in tissue biopsy. Here we found that PCNA protein is present in exosomes in human peripheral blood. Our results demonstrated that PCNA protein is released from cells as a component of exosomal cargoes and can be detected using a variety of immunoassays such as immunoblot and ELISA. We also found that patients diagnosed with cancers generally have higher sera exosomal PCNA protein levels than healthy subjects, suggesting PCNA can be developed as a liquid biopsy molecular marker for cancers. Here, our initial data on its AUROC curve for diagnosing cancers was 0.94 (95%CI: 0.92–0.97) with an ideal sensitivity (77.2%) and specificity (94.4%) under an optimal cut-off value of 97.2ng/ml. Our result demonstrated that PCNA is an excellent serum marker for liquid biopsy that can be used in a variety of oncology laboratory testing applications. Our findings would be a potentially seminal discovery. Further validation studies are needed to confirm the role of PCNA alone or in combination with other markers in the detection and diagnosis of pan-cancers.

PCNA has been used as a tissue-based biomarker of cancers in clinical practice (15, 16). The presence of this cancer-related protein in exosomes is novel. Our results could expand greatly the role of PCNA in oncological applications, such as early detection, treatment monitoring, and prognostic prediction. There are several compelling advantages to serum testing of PCNA. First, exosomal PCNA is secreted from cancer cells to blood long before symptoms appear, which allows early detection of cancer, and promotes better treatment outcomes. Second, as a general proliferation marker, PCNA is highly produced by all malignant cells including the secreted form. Thus, exosomal PCNA can be used as an ideal molecular window for liquid biopsy pan-cancer screening. Early detection is key. Further, screening for multiple cancers in an individual at a time is either too expensive or not tangible, it is desirable to have a multi-cancer screening tool, namely a single test to detect multiple cancers. Our results suggested that PCNA is the best multi-cancer biomarker. We discovered the presence of PCNA in peripheral body fluids and developed methods to detect them. Our research paved the way for developing a new diagnostic tool for oncology that will be more affordable and accessible. Moreover, cancer-associated isoform of PCNA (caPCNA) has been described (17) and may serve as a more effective indicator of malignancies. We are currently studying whether caPCNA can be detected in human peripheral bodily fluids and their potential specific indications.

The discovery of the PCNA protein in the exosomes would also inspire research interest in exploring its non-nuclear functions. For example, cell surface PCNA has been identified as a ligand of the innate immune receptor NKp44 and can enable cancer cell immune evasion through NKp44 receptor inhibiting NK cell attack (22, 23). Cytosolic PCNA has been shown to have roles in controlling metabolism and cell survival in hematological cells (24, 25). Exosomes are effective intercellular communicators (26), therefore, exosomal PCNA may exert its non-nuclear and cell cycle-independent functions in distance organs via circulation.

## Author contributions

Conceptualization: H.L., C.C.; Methodology: W. S., X. S., H. Z., H.L., C.C.; Validation: W. S., X. S., H. Z., H.L., C.C.; Investigation: W. S., X. S., H. Z., H.L., C.C.; Writing-original draft: C.C.; Writing-review and editing: H.L., C.C.; Supervision: H.L., C.C.; Project administration: H.L., C.C.;

## Competing Interests

H.L. and C.C. are co-inventors of a U.S. provisional patent (USPTO patent pending, application #: 63/656,835) which covers the experimental results described here.

## Notes

### Summary of Updates

the affiliations of Chunlin Cai has been changed to correct a writing error

